# Neurophysiological basis of hemodynamic responses in the developing human brain before the time of normal birth

**DOI:** 10.1101/2022.09.23.509234

**Authors:** Tanya Poppe, Jucha Willers Moore, Mohammed Rupawala, Anthony N. Price, Felipe Godinez, Kimberley Whitehead, Sofia Dall’Orso, A. David Edwards, Lorenzo Fabrizi, Tomoki Arichi

**Author notes:** Authors contributed equally to work (Lorenzo Fabrizi, Tomoki Arichi). Corresponding author: Dr Tomoki Arichi, Department of Perinatal Imaging & Health, King’s College London, 1^st^ Floor South Wing, St Thomas’ Hospital, Westminster Bridge Road, London SE1 7EH, UK.

## Abstract

Neurovascular coupling that links neural activity to localized increases in blood flow is essential both for brain function and to prevent tissue injury. In the healthy human brain, this underlies an association between the duration of EEG microstates, which represent coordinated and metastable activation of neuronal ensembles, and increases in hemodynamic activity. However, in early human life it is not clear whether neurovascular coupling is functional as the underlying physiological mechanisms may be too immature to effectively support it. Here, we combined MRI compatible robotics with simultaneous EEG and fMRI data acquisition in 13 preterm infants to assess whether the relationship between neural activity and hemodynamic responses is present in this critical period of early life. Passive sensorimotor stimulation elicited both a distinct sequence of four EEG microstates and a significant rise in the blood oxygen level dependent (BOLD) fMRI signal in the left primary sensorimotor cortex. Furthermore, EEG microstate duration was significantly related to BOLD response amplitude. These results suggest that effective neurovascular coupling is present in the human brain even before the normal time of birth and reveal a complex relationship between EEG and fMRI signals underpinned by patterns of activity across distinct neural ensembles.

## INTRODUCTION

Across the lifespan, the human brain consumes disproportionately high levels of energy relative to its mass (Raichle and Gusnard, 2002). Despite this, the brain lacks a reservoir for the required metabolic substrates and thus critically relies on their rapid provision via the bloodstream (Iadecola, 2017). This is achieved through neurovascular coupling, which cumulatively describes a series of tightly regulated mechanisms that lead to a localized increase in cerebral blood flow within the vasculature supplying active neuronal tissue. This not only enables metabolic demands to be met, but crucially prevents tissue injury and cellular degeneration (Iadecola, 2017). In the healthy mature brain, the presence of neurovascular coupling underpins a direct relationship between electrical neural signals recorded with electroencephalography (EEG) and the hemodynamic response measured with Blood Oxygen Level Dependent (BOLD) functional Magnetic Resonance Imaging (fMRI). A key feature is a significant relationship between the duration of EEG microstates, which represent the coordinated and metastable activation of large neuronal ensembles, and the spatial and temporal features of the fMRI BOLD hemodynamic signal both at rest and when performing tasks (Musso et al., 2010, Britz et al., 2010, Michel and Koenig, 2018).

In the time leading up to full term human birth, rapid maturational changes are taking place across nearly all of the components which both relate to and occur within the neurovascular coupling cascade (Harris et al., 2011). This includes marked developmental increases in: (i) cortical neural density and metabolism; (ii) perivascular cell density and function; (iii) angiogenesis and capillary bed density; (iv) systematic changes in perivascular signaling and (v) global cerebral blood flow and volume (Harris et al., 2011, Kozberg et al., 2013). Together, this could potentially mean that effective neurovascular coupling is not present in the developing brain (Kozberg and Hillman, 2016). However, there is also evidence that sensory stimulation in preterm infants elicits robust, albeit immature, electrophysiological (Fabrizi et al., 2011, Khazaei et al., 2021) and BOLD hemodynamic responses (Allievi et al., 2016, Arichi et al., 2012, Arichi et al., 2010). Understanding the developmental state of neurovascular coupling has clear implications for newborn infants and those in critical care, where sudden and untimely increases in *ex utero* sensory experience on a background of absent or ineffective neurovascular coupling could contribute to the higher rates of brain injury and later adverse neurodevelopmental outcome in this vulnerable population (Brew et al., 2014).

Here we test the hypothesis that despite the apparent immaturity of the underlying physiology, neurovascular coupling is functional before the normal time of birth. To achieve this, we measured induced neural and hemodynamic responses to a sensorimotor stimulus using simultaneous EEG-fMRI acquisition from a group of preterm human infants and assessed whether (i) EEG microstates and fMRI BOLD hemodynamic responses could be concurrently elicited and (ii) EEG microstate features are related to BOLD amplitude.

## RESULTS AND DISCUSSION

Data were successfully acquired from 13 preterm infants without clinically significant brain injury (median gestational age (GA) at birth: 32^+6^, range: 29^+2^ to 35^+1^ weeks^+days^; median post-menstrual age (PMA) at study: 34^+0^, range: 31^+4^ to 36^+0^ weeks^+days^). The sensorimotor stimulus was delivered to their right wrist with a MR-compatible robotic device which elicited safe and reproducible flexion-extension passive movements (*figure 1*) (Dall’Orso et al., 2018, Allievi et al., 2013). In all infants, a positive BOLD hemodynamic response (consistent with a rise in oxygenated hemoglobin and a decrease in deoxygenated hemoglobin) to the stimulus was identified in the contralateral (left) primary sensorimotor cortex, with 9/13 also having a cluster of activity in the ipsilateral (right) hemisphere. No negative BOLD responses were identified. Group analysis showed that the average response was a significant cluster (p<0.05, family wise error corrected) that localized to the left primary sensorimotor cortex (*figure 2*). The median BOLD hemodynamic response peaked at 14 seconds after stimulus onset, with a median BOLD signal change of 0.56%.

**Figure 1:**
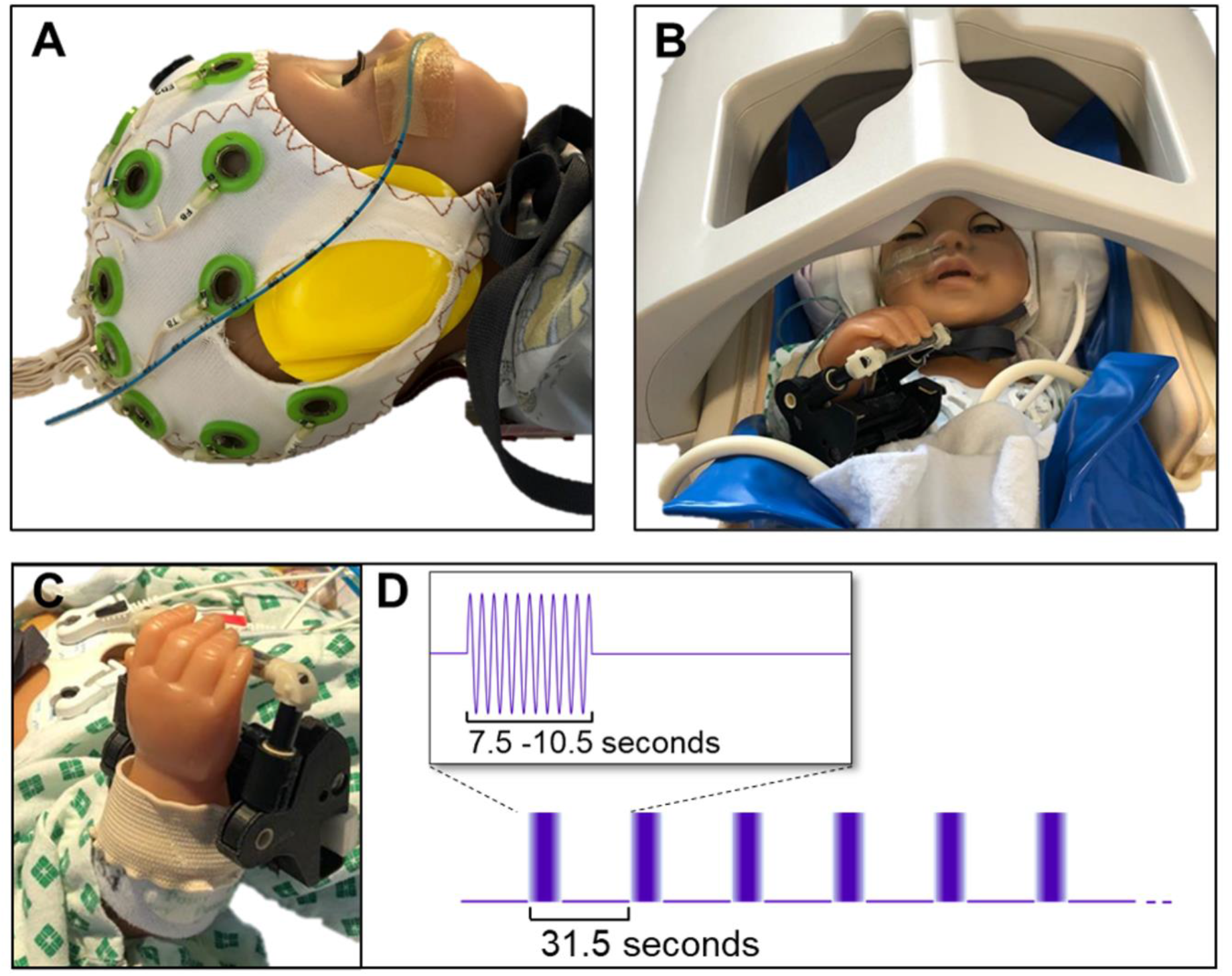
Experimental design involved simultaneous electroencephalography and functional magnetic resonance imaging (EEG-fMRI) data collection from preterm infants whilst receiving passive sensorimotor stimulation. (A) Infants were fitted with a custom MR compatible 25-electrode EEG cap and hearing protection. (B) They were then placed inside the MR head coil, positioned with inflatable head padding (Pearltec, Zurich CH) and a vacuum evacuated immobilization blanket (blue) and data were acquired during natural sleep. (C) To elicit sensorimotor neural activity, a custom-built MR compatible robotic device was fitted to the infants’ right wrist. D) The experiment then consisted of a block design paradigm during which the device passively flexed and extended the right wrist in 1 Hz cycles for 7.5 to 10.5 seconds, a a full block lasted 31.5 seconds.

Data analysis of EEG response epochs (76 epochs of 28.5s) revealed a corresponding rise in mean global field power (GFP) during the stimulus period (*figure 3B*), which had long periods of significant topographic consistency across trials/subjects (not present during the interstimulus interval). These periods could be clustered into four different microstates (*figure 2a*) which accounted for 82% of the total signal variance during the stimulus period and remained stable for significantly longer periods during the stimulus (mean duration: 314ms; standard deviation (SD): 155ms) than in the interstimulus interval (mean duration: 177ms; SD: 155ms) (p=0.03). This result suggests that the engagement of these microstates was stimulus-related and that sensorimotor processing, even at this early stage of life, is considerably more complex than implied by the unilateral focal hemodynamic response identified with fMRI alone. In the mature brain, the sequential engagement of microstates has been found to represent different phases of hierarchical somatosensory processing and/or top-down modulation of sensation driven by different neural populations (Hu et al., 2014). Thus, the observed composite progression of microstates indicates that the preterm brain is already capable of multi-level local sensory elaboration in the primary sensorimotor cortices. This complexity in the sensorimotor response likely reflects relative maturity which is consistent with previous work demonstrating somatotopically organized responses in preterm infants (Dall’Orso et al., 2018) and apparent advanced maturation of the temporal and spatial properties of the primary sensory resting state networks in comparison to the putative higher order associative networks (Fransson et al., 2011). However, further work is needed to elucidate if microstate engagement is present during processing related to other sensory modalities and how this links to brain maturation and behavior.

**Figure 2:**
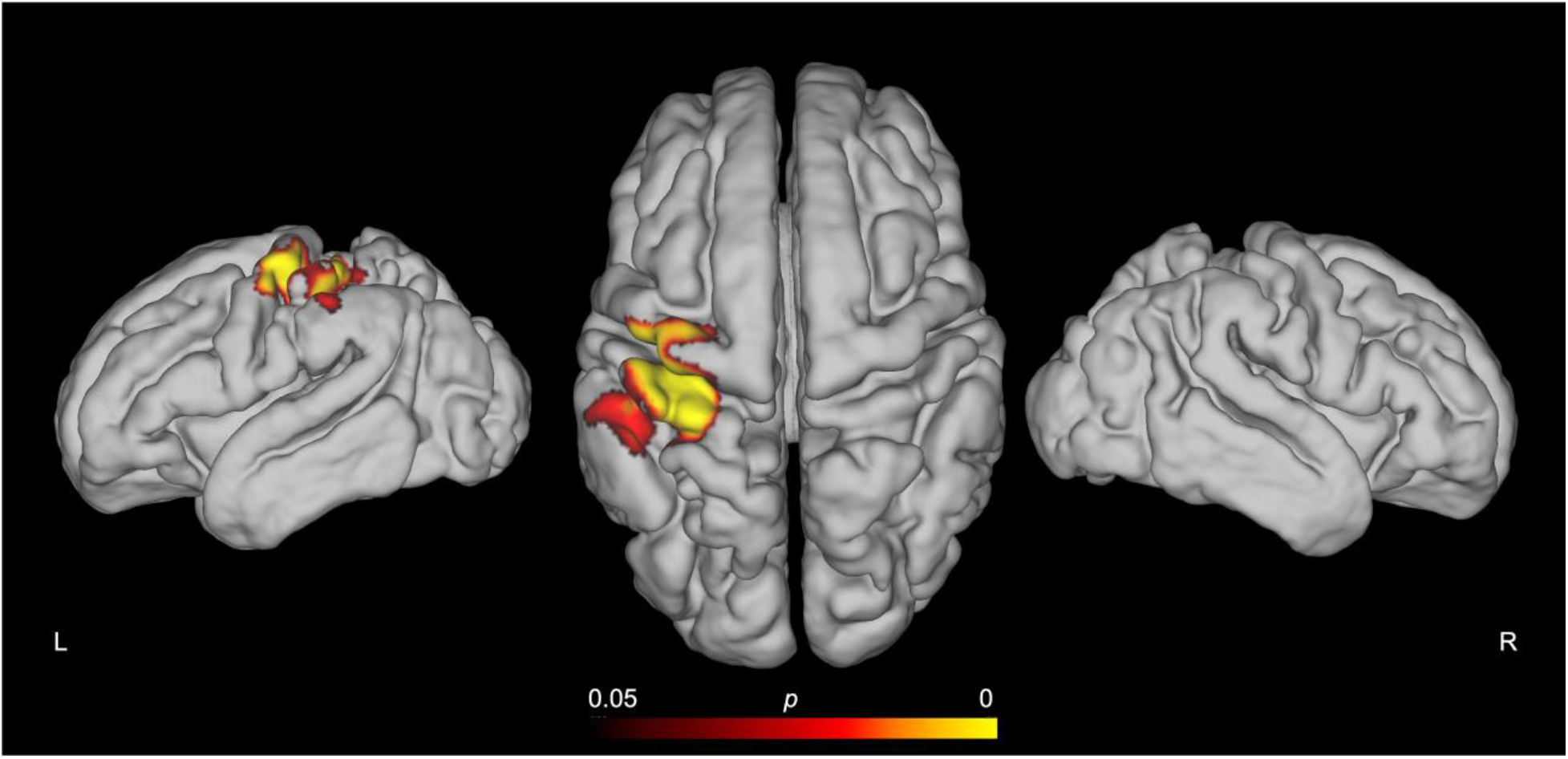
Sensorimotor stimulation induced a localized cluster of positive blood oxygen level dependent (BOLD) activity in the contralateral primary sensorimotor cortex of preterm human infants. Cortical surface projections (left lateral, dorsal, and right lateral) of the group average functional magnetic resonance imaging (fMRI) response to a right wrist flexion-extension passive sensorimotor stimulus from 13 preterm infants of median postmenstrual age 34^+0^ weeks (range 31^+4^ to 36^+0^ weeks). Images show the results of a one-sample nonparametric t-test (*p* < 0.05 corrected for family wise error) projected onto the pial surface of a 34 weeks’ postmenstrual age template brain.

**Figure 3:**
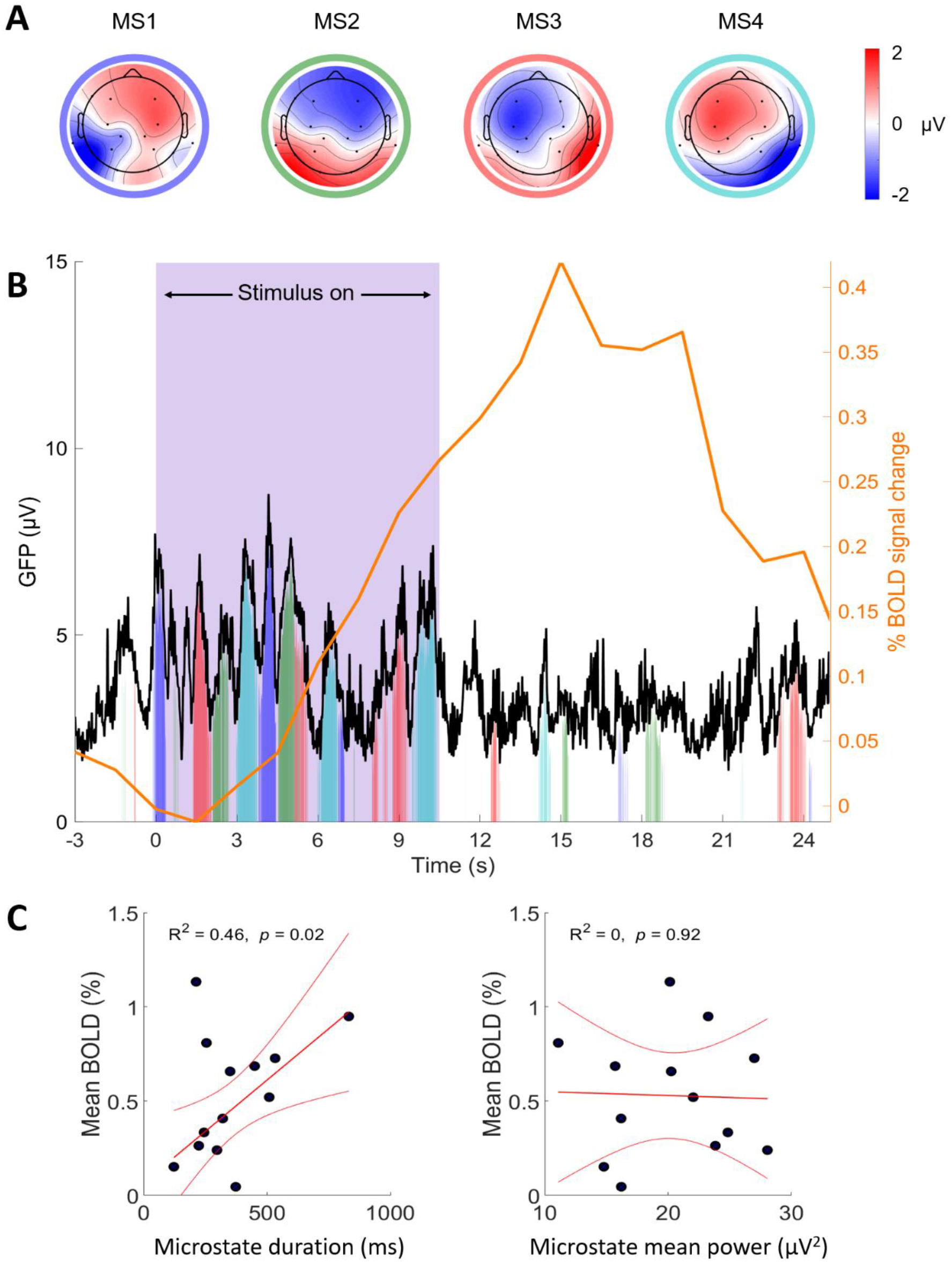
Sensorimotor stimulation induced distinct microstates, the duration of which was significantly related to blood oxygen level dependent (BOLD) response amplitude. **(A)** Topomaps of the four electroencephalogram (EEG) microstates identified from topographic consistency during the stimulus period (n=13, epochs=76). **(B)** Global field power (GFP) of the average EEG signal (black) and the average BOLD signal (orange) across all included stimulus (purple box) epochs, showing a stimulus-related recurrent rise in GFP and a positive BOLD hemodynamic response. Filled color patches represent significant engagement of the microstates in panel (A). The height of the patch represents the amount of signal explained by the microstates at each time-point. **(C)** BOLD response amplitude was significantly related to the total duration of the microstates during the stimulus period (*left:* R^2^=0.46; p<0.02) but not the mean power (*middle:* R^2^ = 0.0; p = 0.92). Black dots represent each subject’s average response, red lines show the estimated linear regression and the 95% confidence intervals.

On a subject level, there was a significant positive correlation between BOLD response amplitude and total microstate duration (R^2^ = 0.46, p = 0.02) during stimulation. In contrast, there was no relationship between the BOLD response amplitude and the infants’ microstate mean power (R^2^ = 0.0; p = 0.92) (*Figure 3c*). This is consistent with adult studies which similarly found that the duration (not power) of intracranial local field potentials is related to BOLD signal magnitude (Murta et al., 2016) and that fMRI activation maps matching canonical resting-state networks can be identified using the onset and duration (but not amplitude) of resting EEG microstates (Musso et al., 2010). In this context, longer microstates likely reflect more stable activation of distinct neural ensembles inducing neurovascular coupling, thereby predicting a stronger BOLD response.

In recent years, fMRI has been widely used to study early human brain development, demonstrating that the preterm period is a key time for the early maturation of resting state brain networks (Doria et al., 2010), the functional connectome (Xu et al., 2019) and organized functional responses (Allievi et al., 2016, Arichi et al., 2012). Crucially, the identified relationship between evoked electrical neural activity and a positive BOLD response not only support that neurovascular coupling is present in this population, but also that the aforementioned fMRI study findings represent true developmental changes in neural processing rather than hemodynamic changes alone. They also highlight the additional insights that can be gained through simultaneous EEG-fMRI data collection, as we demonstrate that apparently simple BOLD responses are instead associated to a multifaceted sequence of different neural states and are therefore only a spatial representation of the complex neuronal activity engaged across the period of stimulation.

Our study is the first exploration of evoked functional responses using simultaneous EEG-fMRI data collection in human infants. Importantly, our findings differ markedly from those from rodent studies where a negative BOLD response without functional hyperemia was described in very young animals, suggesting an absence of neurovascular coupling (Kozberg et al., 2013). They are also not consistent with the proposed relationship between positive BOLD responses and a rise in systemic blood pressure due to a lack of cerebral autoregulation (Kozberg et al., 2013), which would lead to a global increase in cerebral blood flow rather than the localized BOLD responses we observed. Instead, our results, together with our previous work exploring the hemodynamic correlates of spontaneous bursting neural activity in this population (Arichi et al., 2017), support those of studies which have found immature but adult-like positive hemodynamic responses in infants and young animals (Arichi et al., 2012, Colonnese et al., 2008). This suggests that neurovascular coupling matures earlier in humans than in rodent pups, which are commonly considered age-equivalent models in respect to cortical functioning (Khazipov and Luhmann, 2006). To explore and fully understand these differences and their possible role in pathology, studies using comparable methodology are needed to characterize the species-specific maturational trajectories that underlie the establishment of neurovascular coupling in early life. However, our results imply that immature neurovascular coupling may not have a significant role in the pathophysiology of cerebral tissue injuries typically seen in preterm born infants (Volpe, 2009); and even that clinical interventions for perinatal brain injury could account for, accommodate, or capitalise on the presence of neurovascular coupling in the preterm human brain to minimize the severity of the injury and its long-term consequences.

Effective neurovascular coupling as demonstrated by a significant link between EEG microstates and the BOLD hemodynamic response is a fundamental physiological mechanism throughout human life. By demonstrating a significant relationship between coordinated activity across large neuronal ensembles and evoked hemodynamic responses in the preterm human brain, we show that neurovascular coupling is established before the time of normal birth in humans, with clear implications for understanding neural processing and characterizing brain function during this critical phase of early human development in both health and disease.

## ACKNOWLEDGMENTS

We thank the families who generously give their time to take part in medical research during what can be a challenging time in their lives. TP and TA were supported by funding from a Medical Research Council (MRC) UK Clinician Scientist Fellowship [MR/P008712/1] and a Translation Support Award [MR/V036874/1]. JWM is in receipt of PhD funding, ADE and TA received funding support from the MRC Centre for Neurodevelopmental Disorders, King’s College London [MR/N026063/1]. KW was supported by funding from Brain Research UK [201819-23]. LF and MR were supported by an MRC project grant [MR/S003207/1]. The authors additionally acknowledge support from the Department of Health via the National Institute for Health Research (NIHR) comprehensive Biomedical Research Centre award to Guy’s & St Thomas’ NHS Foundation Trust in partnership with King’s College London and King’s College Hospital NHS Foundation Trust, and the Wellcome Engineering and Physical Sciences Research Council (EPSRC) Centre for Medical Engineering at Kings College London [WT 203148/Z/ 16/Z].

## AUTHOR CONTRIBUTIONS

Conceptualization and resources: A.D.E., L.F., T.A.; Methodology: T.P., A.N.P., F.G., K.W., S.D, L.F., T.A.; Data acquisition: T.P., K.W., L.F., T.A.; EEG analysis: T.P., J.W.M, M.R., L.F.; fMRI analysis: T.P., T.A.; All authors reviewed and edited the manuscript.

## METHODS

### Participants

17 preterm infants born at 29^+2^ to 35^+1^ gestational weeks^+days^ and studied on postnatal day 2 to 26, at 31^+4^ to 36^+0^ weeks’ PMA (*Table 1*) were recruited from the postnatal and neonatal intensive care wards at St Thomas’ Hospital, London. Data were later excluded from 4 of the infants due to image artifacts and excessive head motion during data acquisition. Informed parental consent was given prior to all data collection. The research methods conformed to the standards set by the Declaration of Helsinki and were approved by the National Research Ethics Committee (12/LO/1247). All infants were clinically assessed by a pediatrician prior to study and were excluded if they required respiratory support, had a history of clinically significant or severe brain pathology such as intraventricular hemorrhage greater than grade 3, birth asphyxia, focal intracerebral lesions affecting the parenchyma or white matter, severe hydrocephalus, or congenital brain malformations.

**Table 1.** Clinical demographics of all preterm neonates scanned and those included in the analysis.

|  | **Scanned (n=17)** |  | **Included (n=13)** |  |
| --- | --- | --- | --- | --- |
|  | **median** | **(range)** | **median** | **(range)** |
| Gestational age at birth (weeks + days) | 32<sup>+0</sup> | (29<sup>+2</sup> - 35<sup>+1</sup>) | 32<sup>+6</sup> | (29<sup>+2</sup> - 35<sup>+1</sup>) |
| Sex (female/male) | 5/12 |  | 5/8 |  |
| Birthweight (kg) | 1.67 | (1.00 - 2.83) | 1.67 | (1.12 - 2.46) |
| Postmenstrual age at scan (weeks + days) | 34<sup>+0</sup> | (31<sup>+4</sup> - 36<sup>+0</sup>) | 34<sup>+0</sup> | (31<sup>+4</sup> - 36<sup>+0</sup>) |
| Postnatal age at scan (days) | 10 | (2 - 26) | 9 | (2 - 23) |
| Head circumference at scan (cm) | 30 | (26 - 34) | 31 | (28 - 34) |

### EEG-fMRI acquisition

MRI data were acquired with a 3-Tesla Philips Achieva scanner (Best, Netherlands) and a 32 channel adult head coil. Prior to scanning, infants were fitted with neonatal 25-electrode EEG caps (29 - 32 cm head circumference) and connected to an MR-compatible system (EASYCAP and Brain Products GmbH). Infants were scanned following feeding, during natural sleep and were fitted with ear protection (molded dental putty and adhesive earmuffs: Minimuffs, Natus Medical Inc, San Carlos CA, USA) and immobilized in a vacuum cushion (Med-Vac, CFI Medical Solutions, Fenton, MI, USA). fMRI data were acquired using T2*-weighted single-shot gradient echo echo-planar imaging (GRE-EPI) sequence (resolution: 2.5*2.5*3.25mm; 21 slices; TE: 30ms; TR: 1500ms, flip angle: 90°, lasting up to 13.5 minutes). A custom-built MR compatible robotic device (*see figure 1* (Dall’Orso et al., 2018)) was fitted to the right wrist to deliver blocks of 1Hz passive right wrist flexion-extension for 7.5 to 10.5 seconds (5 to 7 TRs, up to 24 epochs of stimulation), with a variable inter-stimulus interval (21 to 24 seconds) to minimize anticipatory responses. High resolution T1 (Magnetization-prepared Rapid Gradient Echo, MPRAGE; resolution: 0.8mm isotropic; TE: 4.6ms; TR: 17ms, flip angle: 13°) and a modified low-specific absorption rate (SAR) T2 (Turbo Spin Echo, TSE; resolution: 0.9mm isotropic; 49 axial slices; TSE factor: 16; TE: 160ms) anatomical scans were also acquired for image registration. All images were formally reported by a neonatal neuroradiologist.

### fMRI pre-processing and analysis

Analysis was performed using tools implemented in FSL (https://fsl.fmrib.ox.ac.uk/fsl/fslwiki). fMRI data were cropped to exclude motion, with a minimum required inclusion for analysis of 3 consecutive stimulus blocks. Vascular and motion-related signal artifact was denoised using independent component analysis (MELODIC v3.0) (Beckmann et al., 2005). Activated brain regions were identified using a general linear model (FEAT v5.6) with stimulus timing convolved with age specific HRF basis functions, and mean percent signal change calculated from the top quartile of significant voxels in the left sensorimotor cluster (Arichi et al., 2012). The group average response from unthresholded individual Z-stat maps was derived using non-parametric permutation methods (Randomise v2.0 (Winkler et al., 2014)), adjusted for gestational age at birth and postmenstrual age at scan.

### EEG pre-processing and analysis

EEG data preprocessing was performed using Analyzer 2 software (Brain Products GmbH), with an initial 0.2 Hz high-pass filter used to remove slow frequency drift in the EEG data. After exclusion of TRs with visible motion on the raw EEG, MR gradient artifact was cleaned using a 25 TR sliding window template subtraction. A 40 Hz lowpass and 50 Hz notch filter were applied. Electrodes with poor signal or bridged to the reference (FCz) were removed. Further analysis was done using EEGLAB (https://sccn.ucsd.edu/eeglab/index.php) and Matlab 2021a (The Mathworks, Natick MA). Up to four electrodes were interpolated to achieve 18 common electrodes for all infants. Data were segmented into 31.5 second (21 TR) epochs (−3s to 28.5s relative to stimulus onset) and epochs with any motion artefact were rejected. Data were aligned to account for stimulus onset jitter using a topographic Woody filter and cropped to the minimum common −3 to 27 seconds (−2 to 18 TRs). A topographic Woody filter is an extension of the traditional Woody filter (Woody, 1967) where the iterative correlation and averaging is not performed on a single channel but is summed across all channels.

Microstate analysis was conducted using Ragu (https://www.thomaskoenig.ch/index.php/work/ragu) and custom-written MATLAB functions. We took a geometric approach to microstate analysis, where microstates were considered as basis vectors of a reduced subspace of the original channel space. Our microstate basis vectors were derived from periods within the grand average signal that were topographically consistent across trials/subjects (Koenig and Melie-Garcia, 2010). These samples were clustered using a modified agglomerative hierarchical clustering algorithm from that provided in the Ragu toolbox with the optimal number of microstates selected as the knee point within a plot of the percentage change in the explained variance as a function of the number of clusters. This point was marked as the last cluster set before an additional aggregation led to an average drop in variance of more than 5%. Once the microstate basis vectors were defined, we calculated their projection on individual trials. We then calculated the total duration and power of microstate occurrences in each trial and the average within each subject. The subject-specific BOLD percent signal change was then regressed against each of these parameters.

## REFERENCES

Allievi, A. G., Arichi, T., Tusor, N., Kimpton, J., Arulkumaran, S., Counsell, S. J., Edwards, A. D. & Burdet, E. 2016. Maturation of Sensori-Motor Functional Responses in the Preterm Brain. Cereb Cortex, 26, 402–413.

Allievi, A. G., Melendez-Calderon, A., Arichi, T., Edwards, A. D. & Burdet, E. 2013. An fMRI compatible wrist robotic interface to study brain development in neonates. Ann Biomed Eng, 41, 1181–92.

Arichi, T., Fagiolo, G., Varela, M., Melendez-Calderon, A., Allievi, A., Merchant, N., Tusor, N., Counsell, S. J., Burdet, E., Beckmann, C. F. & Edwards, A. D. 2012. Development of BOLD signal hemodynamic responses in the human brain. Neuroimage, 63, 663–73.

Arichi, T., Moraux, A., Melendez, A., Doria, V., Groppo, M., Merchant, N., Combs, S., Burdet, E., Larkman, D. J., Counsell, S. J., Beckmann, C. F. & Edwards, A. D. 2010. Somatosensory cortical activation identified by functional MRI in preterm and term infants. Neuroimage, 49, 2063–71.

Arichi, T., Whitehead, K., Barone, G., Pressler, R., Padormo, F., Edwards, A. D. & Fabrizi, L. 2017. Localization of spontaneous bursting neuronal activity in the preterm human brain with simultaneous EEG-fMRI. Elife, 6.

Beckmann, C. F., Deluca, M., Devlin, J. T. & Smith, S. M. 2005. Investigations into resting-state connectivity using independent component analysis. Philos Trans R Soc Lond B Biol Sci, 360, 1001–13.

Brew, N., Walker, D. & Wong, F. Y. 2014. Cerebral vascular regulation and brain injury in preterm infants. Am J Physiol Regul Integr Comp Physiol, 306, R773–86.

Britz, J., Van De Ville, D. & Michel, C. M. 2010. BOLD correlates of EEG topography reveal rapid resting-state network dynamics. Neuroimage, 52, 1162–70.

Colonnese, M. T., Phillips, M. A., Constantine-Paton, M., Kaila, K. & Jasanoff, A. 2008. Development of hemodynamic responses and functional connectivity in rat somatosensory cortex. Nat Neurosci, 11, 72–9.

Dall’Orso, S., Steinweg, J., Allievi, A. G., Edwards, A. D., Burdet, E. & Arichi, T. 2018. Somatotopic Mapping of the Developing Sensorimotor Cortex in the Preterm Human Brain. Cereb Cortex, 28, 2507–2515.

Doria, V., Beckmann, C. F., Arichi, T., Merchant, N., Groppo, M., Turkheimer, F. E., Counsell, S. J., Murgasova, M., Aljabar, P., Nunes, R. G., Larkman, D. J., Rees, G. & Edwards, A. D. 2010. Emergence of resting state networks in the preterm human brain. Proc Natl Acad Sci U S A, 107, 20015–20.

Fabrizi, L., Slater, R., Worley, A., Meek, J., Boyd, S., Olhede, S. & Fitzgerald, M. 2011. A shift in sensory processing that enables the developing human brain to discriminate touch from pain. Curr Biol, 21, 1552–8.

Fransson, P., Aden, U., Blennow, M. & Lagercrantz, H. 2011. The functional architecture of the infant brain as revealed by resting-state fMRI. Cereb Cortex, 21, 145–54.

Harris, J. J., Reynell, C. & Attwell, D. 2011. The physiology of developmental changes in BOLD functional imaging signals. Dev Cogn Neurosci, 1, 199–216.

Hu, L., Valentini, E., Zhang, Z. G., Liang, M. & Iannetti, G. D. 2014. The primary somatosensory cortex contributes to the latest part of the cortical response elicited by nociceptive somatosensory stimuli in humans. Neuroimage, 84, 383–93.

Iadecola, C. 2017. The Neurovascular Unit Coming of Age: A Journey through Neurovascular Coupling in Health and Disease. Neuron, 96, 17–42.

Khazaei, M., Raeisi, K., Croce, P., Tamburro, G., Tokariev, A., Vanhatalo, S., Zappasodi, F. & Comani, S. 2021. Characterization of the Functional Dynamics in the Neonatal Brain during REM and NREM Sleep States by means of Microstate Analysis. Brain Topogr, 34, 555–567.

Khazipov, R. & Luhmann, H. J. 2006. Early patterns of electrical activity in the developing cerebral cortex of humans and rodents. Trends Neurosci, 29, 414–418.

Koenig, T. & Melie-Garcia, L. 2010. A method to determine the presence of averaged event-related fields using randomization tests. Brain Topogr, 23, 233–42.

Kozberg, M. & Hillman, E. 2016. Neurovascular coupling and energy metabolism in the developing brain. Prog Brain Res, 225, 213–42.

Kozberg, M. G., Chen, B. R., Deleo, S. E., Bouchard, M. B. & Hillman, E. M. 2013. Resolving the transition from negative to positive blood oxygen level-dependent responses in the developing brain. Proc Natl Acad Sci U S A, 110, 4380–5.

Michel, C. M. & Koenig, T. 2018. EEG microstates as a tool for studying the temporal dynamics of whole-brain neuronal networks: A review. Neuroimage, 180, 577–593.

Murta, T., Hu, L., Tierney, T. M., Chaudhary, U. J., Walker, M. C., Carmichael, D. W., Figueiredo, P. & Lemieux, L. 2016. A study of the electro-haemodynamic coupling using simultaneously acquired intracranial EEG and fMRI data in humans. Neuroimage, 142, 371–380.

Musso, F., Brinkmeyer, J., Mobascher, A., Warbrick, T. & Winterer, G. 2010. Spontaneous brain activity and EEG microstates. A novel EEG/fMRI analysis approach to explore resting-state networks. Neuroimage, 52, 1149–61.

Raichle, M. E. & Gusnard, D. A. 2002. Appraising the brain’s energy budget. Proc Natl Acad Sci U S A, 99, 10237–9.

Volpe, J. J. 2009. The encephalopathy of prematurity--brain injury and impaired brain development inextricably intertwined. Semin Pediatr Neurol, 16, 167–78.

Winkler, A. M., Ridgway, G. R., Webster, M. A., Smith, S. M. & Nichols, T. E. 2014. Permutation inference for the general linear model. Neuroimage, 92, 381–97.

Woody, C. D. 1967. Characterization of an adaptive filter for the analysis of variable latency neuroelectric signals. Medical and biological engineering, 5, 539–554.

Xu, Y., Cao, M., Liao, X., Xia, M., Wang, X., Jeon, T., Ouyang, M., Chalak, L., Rollins, N., Huang, H. & He, Y. 2019. Development and Emergence of Individual Variability in the Functional Connectivity Architecture of the Preterm Human Brain. Cereb Cortex, 29, 4208–4222.

